# IgA MAb blocks SARS-CoV-2 Spike-ACE2 interaction providing mucosal immunity

**DOI:** 10.1101/2020.05.15.096719

**Authors:** Monir Ejemel, Qi Li, Shurong Hou, Zachary A. Schiller, Aaron L. Wallace, Alla Amcheslavsky, Nese Kurt Yilmaz, Jacqueline R. Toomey, Ryan Schneider, Brianna J. Close, Da-Yuan Chen, Hasahn L. Conway, Saeed Mohsan, Lisa A. Cavacini, Mark S. Klempner, Celia A. Schiffer, Yang Wang

## Abstract

COVID-19 caused by SARS-CoV-2 has become a global pandemic requiring the development of interventions for the prevention or treatment to curtail mortality and morbidity. No vaccine to boost mucosal immunity or as a therapeutic has yet been developed to SARS-CoV-2. In this study we discover and characterize a cross-reactive human IgA monoclonal antibody, MAb362. MAb362 binds to both SARS-CoV and SARS-CoV-2 spike proteins and competitively blocks hACE2 receptor binding, by completely overlapping the hACE2 structural binding epitope. Furthermore, MAb362 IgA neutralizes both pseudotyped SARS-CoV and SARS-CoV-2 in human epithelial cells expressing hACE2. SARS-CoV-2 specific IgA antibodies, such as MAb362, may provide effective immunity against SARS-CoV-2 by inducing mucosal immunity within the respiratory system, a potentially critical feature of an effective vaccine.

## Introduction

In December 2019, a novel coronavirus (SARS-CoV-2) was identified as the cause of an outbreak of acute respiratory infections that emerged in Wuhan, China. The coronavirus disease 2019 (COVID-19) ranges from *mild* to *severe acute respiratory* infection, with a fatality rate estimated *to range* from 2 to *3*%^1-4^. Within three months of the first report cases, COVID-19 rapidly disseminated through the human population and had become a global pandemic by March 2020. Phylogenic analysis has classified SARS-CoV-2 within the sarbecoviruses subgenus, the β lineage that also contains SARS-CoV, sharing proximately 79.6% sequence identity^4^.

Interventions for the prevention or treatment of COVID-19 are crucial for the ongoing outbreak. Pre-or post-exposure immunotherapies with neutralizing antibodies, would be of great use by providing immediate mucosal immunity against SARS-CoV-2. Although concerns, as occurred with SARS-CoV^5,6^, that vaccines may cause disease enhancement will need to be addressed. The feasibility of human monoclonal antibodies (MAbs) as immunoprophylaxis or therapy against coronaviruses including SARS-CoV^7-10^ and MERS-CoV^11^ has been demonstrated. These anti-coronavirus MAbs primarily target the viral spike (S) glycoprotein, a type I transmembrane glycoprotein that produces recognizable crown-like spike structures on the virus surface. The receptor-binding domain (RBD) of the S protein facilitates viral entry into human cells through human angiotensin-converting enzyme 2 (hACE2) receptor binding leveraging a similar mechanism as SARS-CoV^12-14^.

Most current anti-SARS-CoV MAbs neutralize virus by binding to epitopes on the spike protein RBD of SARS-CoV^15^. We and others have demonstrated that neutralizing MAbs that block RBD-hACE2 binding could confer potent protection against SARS-CoV as both prophylaxis and treatment in various animal models^7,9,10^. Several anti-SARS-COV MAbs have demonstrated cross-neutralizing activities against the S protein of SARS-CoV-2^16,17^. However, to date no antibody has directly bound to the hACE2 RBD interface of SARS-CoV-2 competitively blocking the SARS-CoV-2 spike:hACE2 complex.

Antibody-dependent enhancement of viral infections are major *hurdles* in the development of effective vaccines. This enhancement is likely facilitated by the Fc domain of IgG but not for its isotype variant IgA^18^. The avidity of mucosal IgA, in comparison with IgG, due to the multimeric structure, enhances the antibody binding with antigens. In addition, the diverse, high level of glycosylation of IgA antibodies, further protects the mucosal surface with non-specific interference. In animal models, high titers of mucosal IgA in the lung is correlated with reduced pathology upon viral challenge with SARS-CoV^19^. How precisely which isotype may protect the mucosa from SARS-CoV-2 infection remains an open question.

In the current study we describe the discovery of a cross-neutralizing human IgA monoclonal antibody, MAb362 IgA. This IgA antibody binds to SARS-CoV-2 RBD with high affinity directly competing at the hACE2 binding interface by blocking interactions with the receptor. MAb362 IgA neutralizes both pseudotyped SARS-CoV and SARS-CoV-2 in human epithelial cells expressing ACE2. Our results demonstrate that IgA isotype, plays a critical role in SARS-CoV-2 neutralization.

## Results

### Selection of MAb binding to RBD of SARS-CoV-2 in ELISA

We have previously developed and characterized a panel of human MAbs that targets the RBD of the SARS-CoV S glycoprotein, isolated from transgenic mice expressing human immunoglobulin genes^9,10^. These 36 hybridomas were recently screened against the SARS-CoV-2 Spike protein for potential cross-bind activity. MAb362 was identified with cross-binding activity against both the S1 subunit of the SARS-CoV S_1-590_ and SAR-CoV-2 S_1-604_ proteins (**Extended Data Table 1).**

While both IgG and IgA are expressed at the mucosa, IgA is more effective on a molar basis and thus the natural choice for mucosal passive immunization as we recently demonstrated in other mucosal infectious disease^20,21^. To further characterize the functionality of MAb362, variable sequences of MAb362 were cloned into expression vectors as either IgGor IgA1 isotypes. Both MAb362 IgG and IgA were assessed in ELISA binding assays against the receptor-binding domain (RBD) of the S1 subunit for SARS-CoV S_270-510_ and SARS-CoV-2 S_319-541_ (**Figure 1a**). MAb362 IgA showed better binding activities, compared to its IgG counterpart against with SARS-CoV-2 S_319-541_ (**Figure 1b**). Assessment of the binding kinetics was consistent with the ELISA binding trends, the binding affinity of IgA with RBD of SARS-CoV-2 is significantly higher (0.3 nM) than that of IgG (13 nM) due to a much slower dissociation rate as an IgA (K_off_ = 1.13×10^−3^ ± 1.06×10^−4^) compared to an IgG (K_off_ = 7.75×10^−5^ ± 5.46×10^−5^) (**Figure 1c-f**).

**Figure 1.**
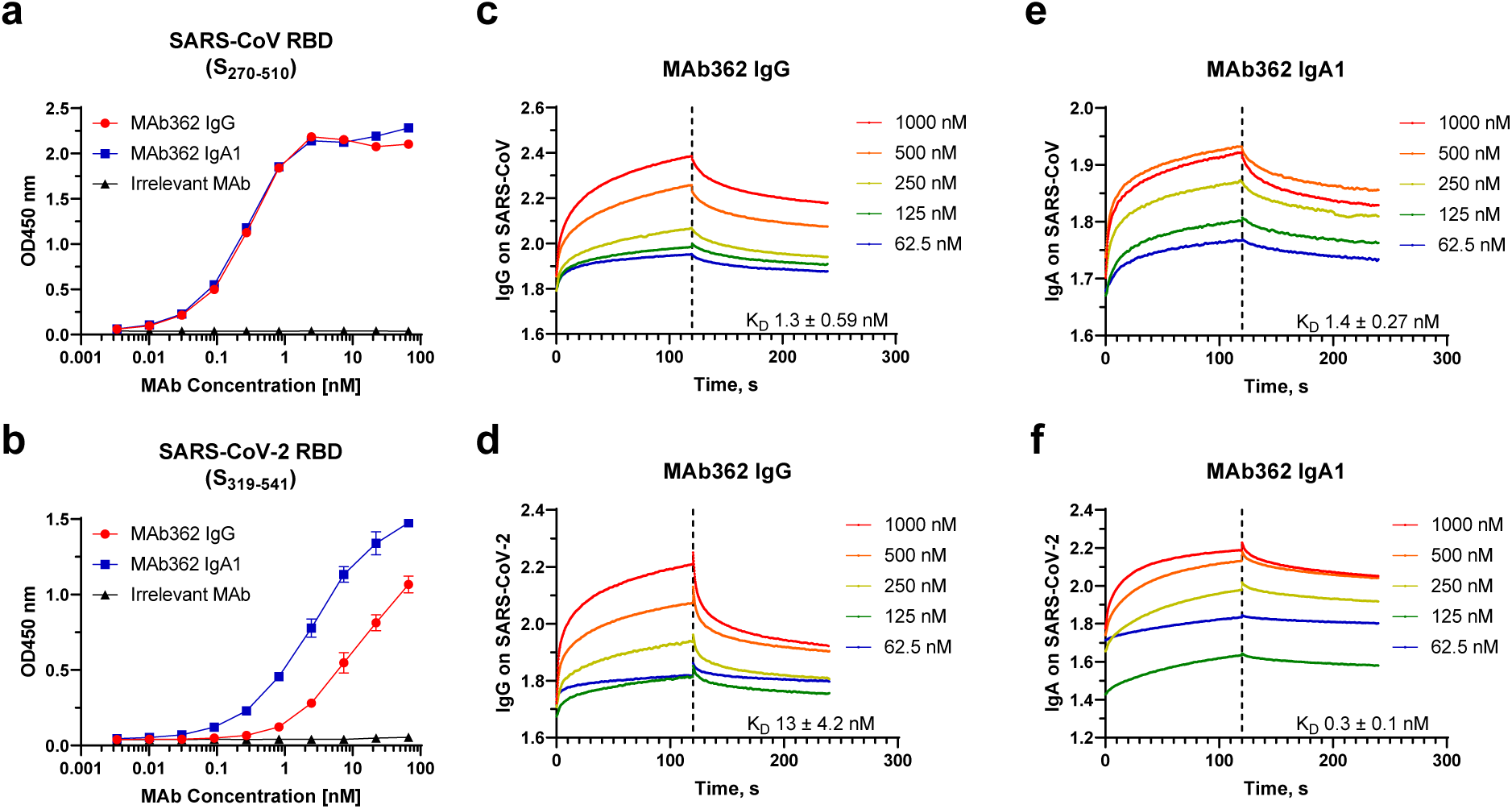
Binding of MAb362 IgG and IgA1 to the RBD of SARS-CoV and SARS-CoV-2. MAb362 IgG and IgA1 bind to purified RBD truncations of the S glycoprotein of SARS-CoV (S_270-510_) (**a**) and SARS-CoV-2 (S_319-541_) (**b**). Affinity measurements of MAb362 IgG (**c, d**) and IgA1 (**e, f**) against the RBD truncations of S glycoprotein of SARS-CoV (S_270-510_) and SARS-CoV-2 (S_319-541_) were conducted using bio-layer interferometry and demonstrate nano and sub-nanomolar affinities. Data is plotted as the average ± SD from two independent experiments.

### Structural modeling MAb362 binding to the core domain of RBD and competes for hACE2 binding

To define the antibody-binding epitope, known co-crystal and cryo-EM complexes from SARS-CoV and MERS spike protein in complex with neutralizing antibodies were evaluated for their potential to competitively block hACE2 binding, based on the structural interface of hACE2-SARS-CoV-2-RBD (PDB ID-6VW1)^22^. The 80R-SARS-CoV-RBD complex (PDB ID-2GHW) ^23^, a crystal structure of SARS-CoV RBD in complex with a neutralizing antibody, *80R* was found most closely to have these characteristics. When the sequence was evaluated, we ascertained that the two antibodies, MAb362 and 80R had frameworks with *striking 90*% amino acid *sequence identity*. Thus, the crystal structure 2GHW provided an outstanding scaffold to build a highly accurate atomic homology model of MAb362. This structure permitted the modeling with the superposition of the hACE2:SARS-CoV-2-RBD (PDB ID-6VWI) for the modeling of MAb362:SARS-CoV-2-RBD (**Figure 2a**).

**Figure 2.**
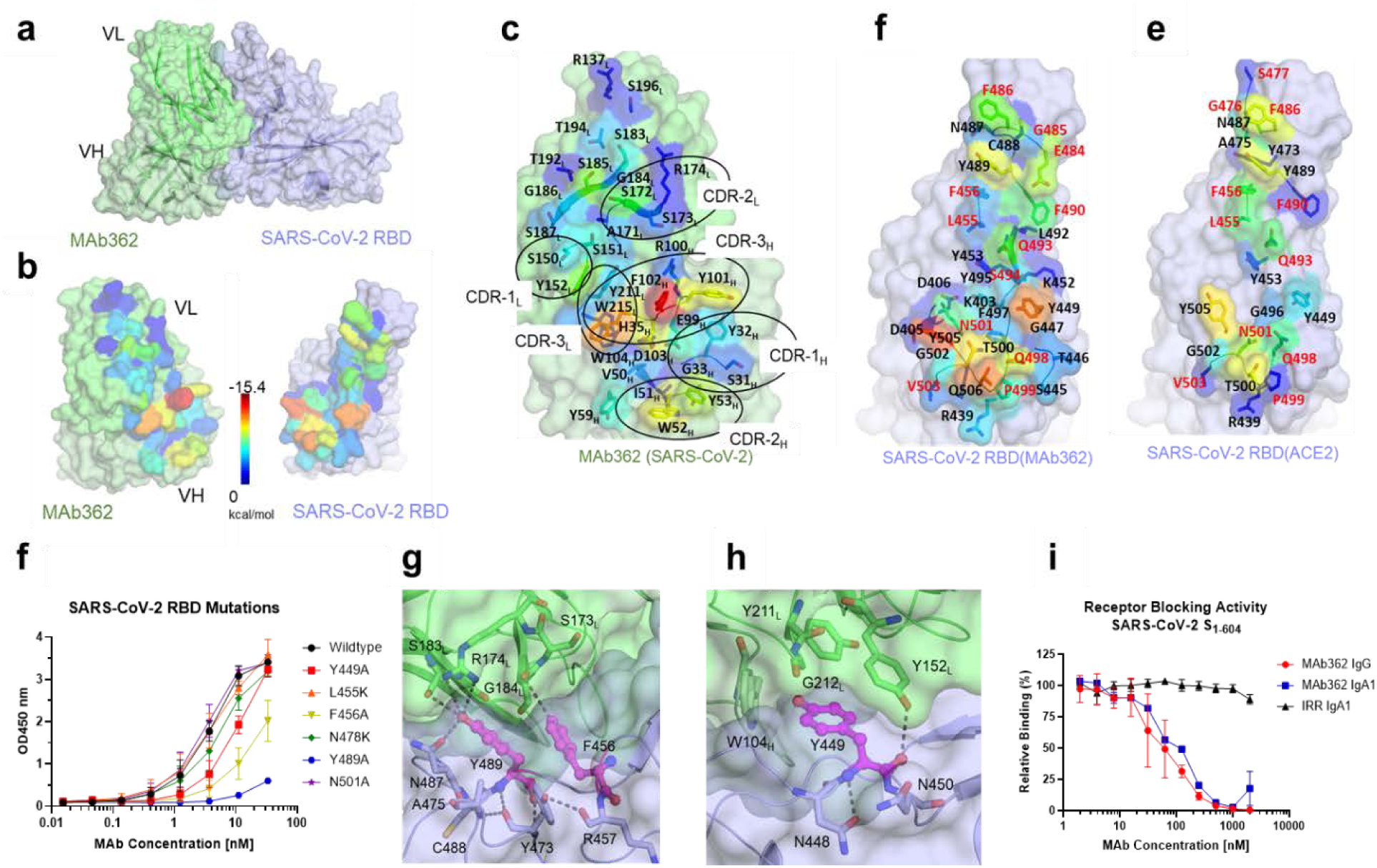
Structural modeling MAb362 binding to the core domain of RBD and competes for hACE2 binding. MAb362 is predicted to have an extensive complementary interface with SARS-CoV-2 RBD that extensively overlaps with the hACE2 binding interface. (**a**) Surface representation of the complex, MAb362 in green, with SARS-CoV-2 RBD in violet. **(b)** The complex is separated and rotated by ∼45° to highlight the extensive van der Waals buried surface areas on each protein, the spectrum of color coding represents the extent of the predicted van der Wall contact, with red being the most extensive contact and dark blue the least. **(c)** Detailed van der Waals of MAb362, residues from all the CDR’s from both heavy and light chains pack against the SARS-CoV-2 RBD. **(d)** The binding interface on SARS-CoV-2 RBD with MAb362. **(e)** The binding interface on SARS-CoV-2 RBD with hACE2. Residues names shown in red are those that differ in sequence between SARS-CoV and SARS-CoV-2. (**f**) Binding affinity of wildtype and six SARS-CoV-2 RBD variants validated critical residues identified by SARS-CoV-2 RBD modeling. Data is plotted as the average ± SD from three independent experiments. (**g**) MAb362 IgG and IgA1 dose dependent reduction of S_1_ truncation of SARS-CoV-2 (S_1-604_) binding to hACE2 receptors on transfected Vero E6 cells. Data is plotted as the average ± SD from two independent experiments. (**h**) Interactions of SARS-CoV-2 RBD (violet) Y489 and F456 (in pink) with MAb362 (green) dotted lines indicate potential hydrogen bonds. (**i**) Interactions of SARS-CoV-2 RBD (violet) Y449 (pink) with MAb362 (green).

The interface between the MAb362:SARS-CoV-2-RBD complex is predicted to form extensive van der Waals contacts (**Figure 2b**). The CDRs of both the heavy and light chain make extensive interactions with SARS-CoV-2-RBD (**Figure 2c**), with the heavy chain of CDR-3 having the most extensive interaction. The binding interface of MAb362 is predicted to form 32 extensive contacts with residues on SARS-CoV-2-RBD (12 of which vary in sequence relative to SARS-CoV-RBD shown in red font) (**Figure 2d**). Seventeen of these contacts also are major points of contact between hACE2 on the SARS-CoV-2-RBD (**Figure 2e**). Thus MAb362 appears to directly compete for SARS-CoV-2 binding with hACE2.

Point mutations were engineered into the SARS-CoV-2-RBD based on this model and the overlap with the hACE2-RBD binding interface to further validate this model (**Figure 2f**). A combination of alanine and lysine mutations showed that charge mutations at the periphery (L455K) or outside the interface (N478K) had no impact, while sites that formed more extensive interactions Y449A, F456A and Y489A caused dramatic loss of binding affinity, only N501A retained affinity in a way that suggest a water mediated interaction may preserve this site. Both Y449 and Y489 are conserved with SARS-CoV while F456 is a Leucine (**Extended Data Figure 1-3**). Interestingly examination of the complex structure shows the close stacking of F456 against Y489 (**Figure 2g**) that together forming a combined extensive interface with light chain of MAb362, specifically the hydroxyl Y489 forms both extensive van der Waals interactions and hydrogen network at the interface. F456 forms less direct interactions, but structurally stabilizes the interactions of Y489, which explains the strong impact of the two alanine mutations F456A and Y489A. Y449A also forms extensive interactions (**Figures 2h**). Thus the loss of binding interaction from these site mutation validates our model of this complex as being biologically relevant complex

Complementing the mutational analysis, to correlate the epitope binding with functionality, MAb362 IgG and IgA were tested in a receptor-blocking assay with hACE2 expressing Vero E6 cells. The result suggested that both MAb362 IgG and IgA block SARS-CoV-2 RBD binding to receptors in a concentration dependent manner starting at ∼ 4 nM (**Figure 2i, Extended Data Figure 4**). Thus confirming that the MAb362 epitope is directly competing for the hACE2 binding epitope on SARS-CoV-2 Spike.

**Figure 3.**
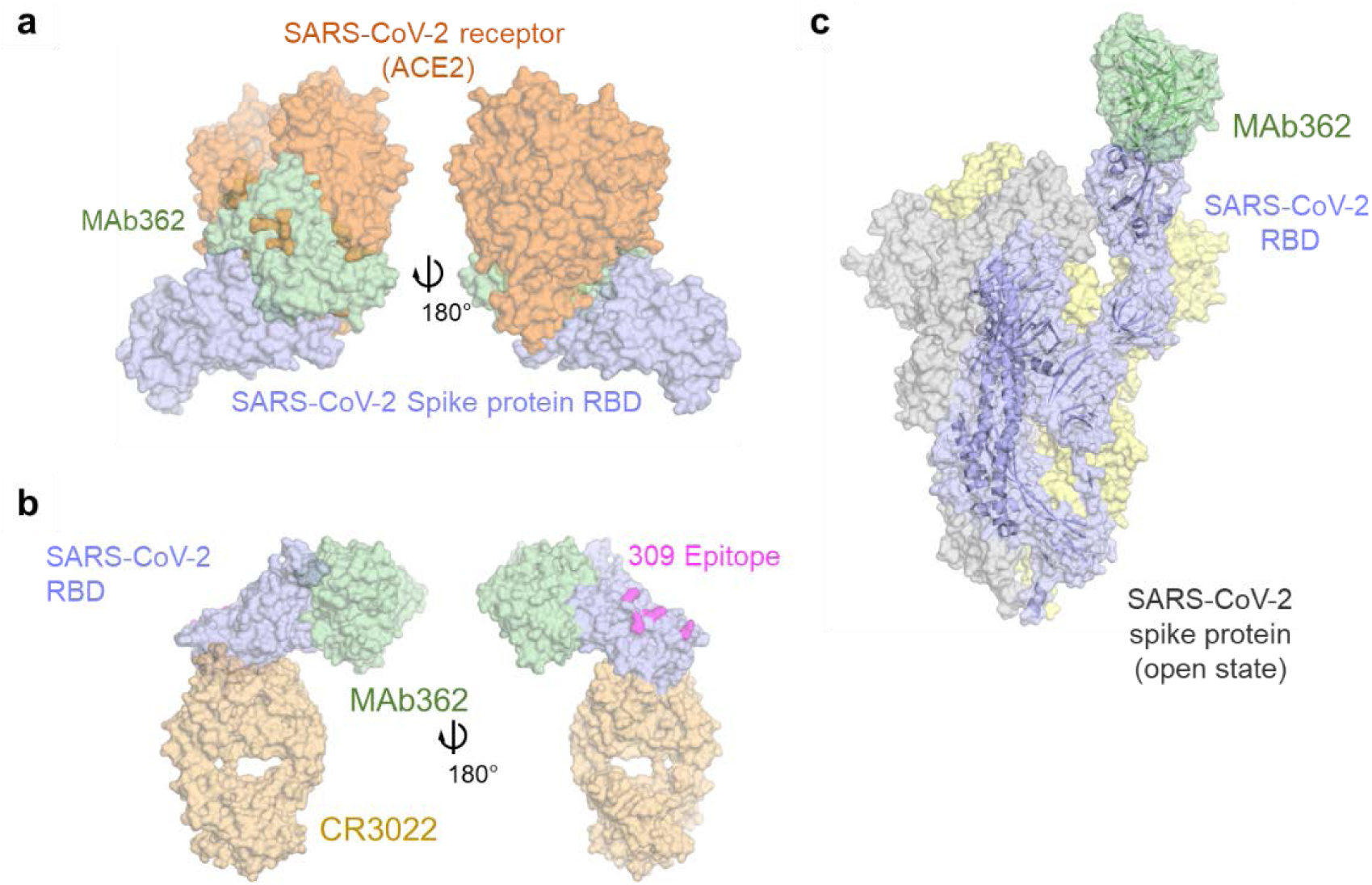
MAb362 structural epitope. (**a**) Superposition of the space filling structures of MAb362 (green) on the complex of hACE2 (orange) -SARS-CoV-2 RBD (violet) (**6VW1)** two views are rotated 180°. (**b**) Positioning of MAb362 on SARS-CoV-2 RBD (violet) relative to the binding of MAbs CR3022^16^ (cream) or 309^17^ epitope (pink). (**c**) MAb362 modeled on the Spike trimer in open conformation with one RBD domain exposed **6VYB**^30^. Note that the binding has no steric clashes with the other monomers.

**Figure 4.**
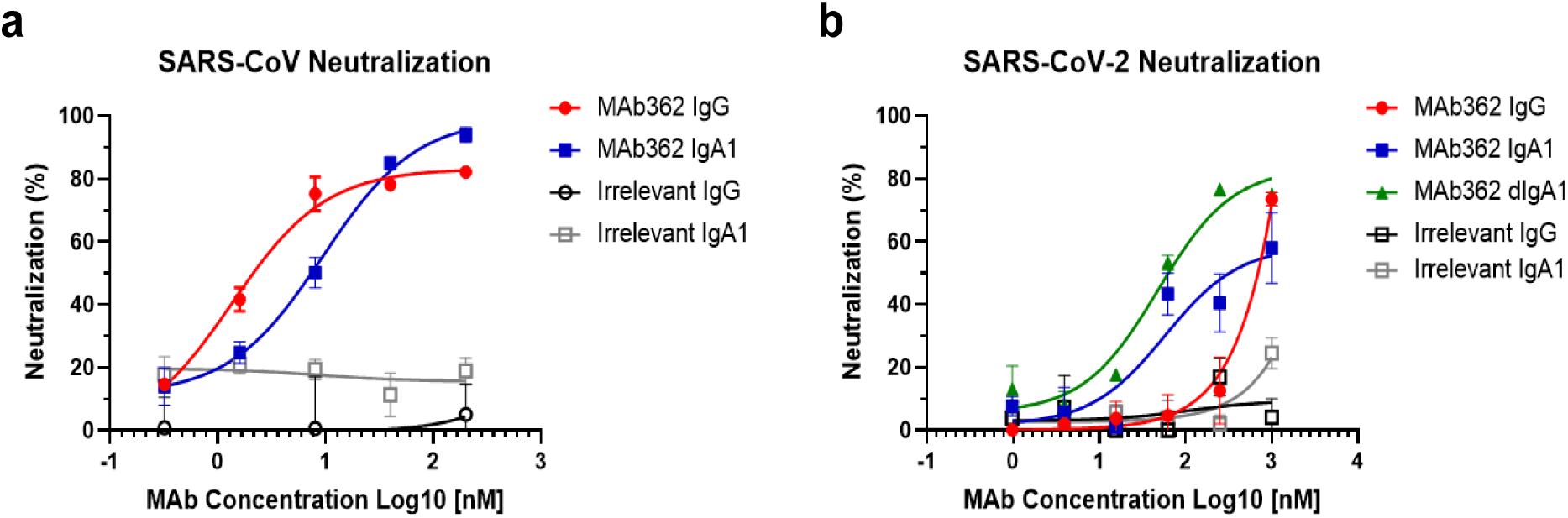
IgA isotype switch enhances MAb362 neutralization of SARS-CoV-2. MAb362 antibody-mediated neutralization of luciferase-encoding pseudovirions with spike proteins of SARS-CoV (**a**) and SARS-CoV-2 (**b**). SARS-CoV and SARS-CoV-2 pseudovirions pre-incubated with fivefold dilutions of MAb362 (0.32 to 200 nM) were used to infect 293 cells expressing hACE2 receptor. Pseudoviral transduction was measured by luciferase activities in cell lysates 48 hrs post transduction to calculate neutralization (%) relative to non-antibody-treated controls. Data is plotted as the average ± SD from two independent experiments.

### MAb362 structural epitope

The epitope of MAb362 is in fact very different from the other recently reported MAb complexes to the SARS-CoV-2-RBD, such as CR3022^16^ or 309^17^. MAb362 overlaps entirely with the hACE2 epitope on the RBD (**Figure 3a**). This contrasts with CR3022 and 309 that bind to epitopes further way from the receptor-binding interface (**Figure 3b**). This finding was consistent with the unique activity of MAb362 of compromising RBD-receptor interaction. As with the binding of hACE2, the MAb362 binding epitope can only be exposed if the RBD was in the open or up conformation in the trimer (**Figure 3c**). In the closed conformation, this epitope would not be accessible to MAb362 without major steric clashes. However, unlike CR3022, MAb362 could access the hACE2 binding epitope(s) if one or more of the trimers is in this open conformation, potentially accounting for the added neutralizing activity.

### MAb362 IgA1 neutralizes SARS-CoV and SARS-CoV-2 better than IgG

To evaluate the neutralization potency of cross-reactive MAb362, we performed a pseudovirus assay using lentiviral pseudovirions on 293 cells expressing hACE2 receptor^24^. Both MAb362 IgG and IgA showed potent neutralization activity against SARS-CoV (**Figure 4a**). MAb362 IgG weakly neutralized SARS-CoV-2 pseudovirus despite its activities to block receptor binding. Interestingly, isotype switch to MAb362 IgA1 resulted in significantly enhanced neutralization potency, compared to its IgG subclass variant (**Figure 4b**). Monomeric MAb362 IgA1 was also co-expressed with J chain to produce dimeric IgA, which further improved neutralization (**Figure 4b**). This is consistent with our prior study showing isotype switch to IgA1 lead to improved antibody neutralization of HIV infection^25^ Our data extends this observation to coronavirus, suggesting that IgA may play an important role in SARS-CoV-2 neutralization.

## Discussion

This study is the first report of a cross-reactive epitope within the core receptor-binding interface of the S protein of both SARS-CoV and SARS-CoV-2. MAb362 IgA neutralizes the virus via directly competing S protein binding to hACE2 receptors. Interestingly, our results show that despite the same blocking of spike interaction with hACE2, MAb362 IgG weakly neutralizes SARS-CoV-2 while its IgA1 isotype variant and its dimeric form showed significantly enhanced neutralization potency. Crystal structure studies demonstrated that IgA1 has a lengthy hinge region with a 13-a.a. insertion and a relaxed “T” like structure as compared to the more rigid “Y” like structure in IgG^26,27^. Thus, the increase flexibility of IgA1 would likely afford a greater reach towards its epitopes on the target and decrease steric hindrance. MAb362 IgA binds when Spike protein (trimer) is in open form. The longer IgA1 hinge may allow two Fabs to reach two RBDs of the trimer at the same time without clashes, which may not be achieved by the shorter hinge in IgG. Our results suggest that compared to IgG, SARS-CoV-2-specific IgA antibody may play an important independent role in providing protective mucosal immunity.

Other recent structure studies have characterized antibodies targeting the RBD domain but distal from the receptor binding core interface of SARS-CoV-2, thus lack the characteristics of how MAb362 blocks the hACE2 binding epitope. Furthermore, these neutralizing IgGs, 47D11 and 309, neutralize SARS-CoV-2 with high potency, but do not block receptor binding to hACE2^17,28^. Potentially hACE2 may not be the sole receptor for SARS-CoV-2, similar to SARS-CoV^29^, or these antibodies may prevent a conformational change necessary for viral entry.

Further study of the interaction between MAb362, and other receptor blocking and neutralizing antibodies against SARS-CoV-2 will provide insight into the design of vaccine and prophylactic/therapeutic antibodies against future emerging infections caused by this viral family.

## Acknowledgements

We thank Dr. Julia Tree at Medical Interventions Group, National Infection Service at Public Health England for support and advice on antibody screen. We thank Dr. Robert Finberg at UMMS for helpful advice on functional assays. The initial mouse immunization work in 2005 was supported by NIH Contract NO1-AI-65315. S.H., N.Y.K. and C.A.S. were supported by NIH R01AI150478

## Author Contributions

M.E., Q.L. cloned, expressed and purified MAbs, S proteins, truncations, and variants with assistance from A.W., A.A., J.T., and R.S. M.E. and Q.L. designed and preformed affinity and binding assays and flow cytometry with assistance from Y.W., L.A.C., and Z.A.S. S.H., carried out structural modeling and analyses with assistance from Q.L. N.Y.K. Y.W. and C.A.S. M.E. and Q.L conducted pseudovirus neutralization with assistance from B.J.C., D-Y.C., H.L.C., S.M., L.A.C., and Y.W. Z.A.S. conducted data and statistical analysis with assistance from M.E., Q.L, and Y.W. Z.A.S. and Y.W. wrote the paper with assistance from M.E., Q.L., L.A.C., M.S.K., and C.A.S. M.S.K., L.A.C., and Y.W. supervised the project.

## Competing interests

A patent application has been filed on 5 May 2020 on monoclonal antibodies targeting SARS-CoV-2 (U.S. Patent and Trademark Office patent application no. 63/020,483; patent applicants: Y.W., M.E., Q.L., and M.K., University of Massachusetts Medical School).

## Materials and Methods

### S glycoprotein expression and purification

SARS-CoV and SARS-CoV-2 S glycoproteins were expressed and purified as previously described^9^. Briefly, the amino acid sequence of the SARS-CoV S glycoprotein (Urbani strain, National Center for Biotechnology Information [strain no. AAP13441]) and SARS-CoV-2 S glycoprotein sequence (GeneBank: MN908947) were used to design a codon-optimized version of the gene encoding the ectodomain of the S glycoproteins a.a. 1–1255 [S_1-1190_] for SARS-CoV and a.a. 1-1273 [S_1-1255_] for SARS-CoV-2, as described elsewhere^31^. The synthetic gene was cloned into pcDNA3.1 Myc/His in-frame with c-Myc and 6-histidine epitope tags that enabled detection and purification. Truncated soluble S glycoproteins were generated by polymerase chain reaction (PCR) amplification of the desired fragments from the vectors encoding S_1255_ and S_1273_. The SARS-CoV-2 RBD constructs carrying point mutation were generated by following the standard protocol from QuikChange® II XL Kit (Agilent). The cloned genes were sequenced to confirm that no errors had accumulated during the PCR process. All constructs were transfected into Expi293 cells using ExpiFectamine(tm) 293 Transfection Kit (Thermo Fisher). Antibodies were purified by immobilized metal chelate affinity chromatography using nickel-nitrilotriacetic acid (Ni-NTA) agarose beads. Antibodies were eluted from the columns using 250 mmol/L imidazole and then dialyzed into phosphate buffered saline (PBS), pH 7.2.

### Generation of MAbs

Previously generated frozen hybridomas of anti-SARS-CoV MAbs^9^ were recovered and scaled up. Hybridoma supernatants were screened for reactivity to the SARS-CoV-2 S protein. Positive cell clones were selected for antibody sequencing. The heavy chain and light chain variable regions were amplified from hybridoma cells and cloned into an immunoglobulin G1 (IgG) expression vector. Isotype switching was conducted using primers designed to amplify the variable heavy chain of the IgG antibody. Products were digested and ligated into a pcDNA 3.1 vector containing the heavy constant IgA1 chain. The vector was transformed in NEB5-α competent cells, and sequences were verified ahead of transient transfection. IgG and IgA1 antibodies were transfected in Expi293 cells and purified as previously described^32^. For dimeric IgA (dIgA), the heavy and light chain vectors were cotransfected with pcDNA-containing DNA for the connecting J chain. Purified antibodies were dialyzed against PBS and then concentrated and quality tested by SDS-PAGE.

### ELISA

Dilutions of purified MAbs were tested in ELISA for reactivity against recombinant S protein. Briefly, 96-well plates were coated with S proteins followed by incubation overnight at 4°C. The plates were blocked with 1% BSA with 0.05% Tween 20 in PBS. Hybridoma supernatant or purified antibody diluted in 1× PBS plus 0.1% Tween 20 and added to the 96-well plates and incubated for 1 hour at room temperature. The plates were stained with horseradish peroxidase-conjugated anti-kappa (1:2,000) for 1 h and developed using 3,3′,5,5′-tetramethylbenzidine. Absorbance at an optical density at 450 nm (OD450) was measured on an Emax precision plate reader (Molecular Devices) using Softmax software.

### Flow cytometry-based receptor binding inhibition assay

Vero E6 cells were harvested with PBS containing 5 mM EDTA and aliquoted to 1×10^6^ cells per reaction. Cells were pelleted then resuspended in PBS containing 10% FBS. Before mixing with the cells, Myc-tagged SARS-CoV S_1-590_ or SARS-CoV-2 S_1-604_ was incubated with the MAb at varying concentrations for 1 hour at room temperature, then the S protein was added to the Vero cells to a final concentration of 10 nM. The cells-S protein mixture was incubated for 1 h at room temperature. After incubation, the cell pellets were washed and then resuspened in PBS with 2% FBS and incubated with 10 µg/mL of anti-Myc antibody for 1 hour at 4°C. Pellets were washed again then subsequently incubated with a Phycoerythrin-conjugated anti-mouse IgG (Jackson Immuno Research) for 40 minutes at 4°C. Cells were washed twice then subjected to flow cytometric analysis using a MACSquant Flow Cytometer (Miltenyi Biotec) and analyzed by FlowJo (version 10). Binding was expressed as relative to cells incubated with S proteins only.

### Pseudotyped virus neutralization assay

Production of pseudotyped SARS-CoV and SARS-CoV-2 was performed as previously described^33^. Pseudovirus was generated employing an HIV backbone that contained a mutation to prevent HIV envelope glycoprotein expression and a luciferase gene to direct luciferase expression in target cells (pNL4-3.Luc.R–E–, obtained from Dr. Nathaniel Landau, NIH). SARS-S and SARS2-S spike protein was provided in *trans* by co-transfection of 293T cells with pcDNA-G with pNL4-3.Luc.R–E–. Supernatant containing virus particles was harvested 48–72 h post-transfection, concentrated using Centricon 70 concentrators, aliquoted and stored frozen at - 80 degree. Before assessing antibody neutralization, the 293T cells were transient transfected with 100ng pcDNA-hACE2 each well in 96 well plates, and the cells were used for the pseudovirus infection 24 hours after transfection. A titration of pseudovirus was performed on 293T cells transiently transfected with hACE2 receptor to determine the volume of virus need to generate 50,000 counts per second (cps) in the infection assay. The appropriate volume of pseudovirus was pre-incubated with varying concentrations of MAbs for 1 h at room temperature before adding to 293T cells expressing hACE2. 24 hours after the infection, the pseudovirus was replaced by the fresh complete media, and 24 hours after media changing the infection was quantified by luciferase detection with BrightGlo luciferase assay (Promega) and read in a Victor3 plate reader (Perkin Elmer) for light production.

### Structural modeling and analyses

Three crystal structures, 2GHW the complex of 80R:SARS-CoV-RBD^23^, 2AJF the complex of hACE2:SARS-CoV-RBD^34^ and 6VW1 the complex of hACE2:SARS-CoV-2-RBD^22^ were used as initial scaffolds in the determinations of the models of MAb362:SARS-CoV-RBD and MAb362:SARS-CoV-2-RBD. The amino acid sequence of MAb362 was aligned to the amino acid sequences of 80R. The molecular modeling of MAb362 was performed through the program Modeller 9.15 using the basic modeling and forming the initial MAb362:SARS-CoV RBD complex. This structure was further refined using iterative energy minimization by Desmond as previously described ^35,36^. The MAb362:SARS-CoV-2-RBD complex was made by replacing the SARS-CoV-2-RBD from 2AJF on the SARS-CoV-RBD structure and further optimized. The structural model of MAb362 binds to the SARS-CoV-2 Spike trimer was based on 6VYB^30^. All figures were made within the PyMOL package. Hydrogen bonds were determined for pairs of eligible donor/acceptor atoms using criteria set by Schrodinger (Schrodinger, LLC, The PyMOL Molecular Graphics System, Version 1.3r1. 2010.). The residue van der Waals potential between the various complexes was extracted from the structures energies using the energy potential within Desmond. Epitope residues predicted by modeling were individually mutated using BioXp(tm) 3200 System (SGI-DNA). The genes were cloned into RBD expression vectors and RBD proteins were purified as described above. Mutant RBD were confirmed intact expression on proteins gels, and the same amount of proteins were coated on the plate for ELISA assays. ELISAs assay was performed to determine binding of the MAbs to the mutant proteins compared to the wild-type.

### Affinity determination

Bio-layer interferometry (BLI) with an Octet HTX (PALL/ForteBio) was used to determine the affinity of MAb362 IgG and IgA1 to the RBD of SARS-CoV and SARS-CoV-2 S protein. MAbs were added to 96 wells plates at 1000 nM and titrated 1:2 to 62 nM using PBS. RBD from SARS-CoV and SARS-CoV-2 were biotinylated (Thermo Fisher) and immobilized on Streptavidin (SA) Biosensors (ForteBio) for 120 seconds at 1600nM concentration. After a baseline step, MAb362-RBD binding rate was determined when the biosensors with immobilized RBD were exposed to MAb362 IgG or IgA1 at different concentrations for 120 seconds. Following association, the MAb362-RBD complex was exposed to PBS and the rate of the MAb362 dissociation from RBD was measured. Each assay was performed in triplicate. c

### Statistical Analysis

Statistical calculations were performed using Prism version 7.03 (GraphPad Software, La Jolla, CA).

