## Supplemental Table and Figures for "IgA MAb blocks SARS-CoV-2 Spike-ACE2 interaction providing mucosal immunity"

| **Clone** | **Epitope**  **(A.A.)** | **SARS-CoV S1 S_1-509_** | **SARS-CoV RBD S_270-510_** | **SARS-CoV-2 S1 S_1-604_** | **SARS-CoV-2 RBD S_319-541_** |
| --- | --- | --- | --- | --- | --- |
| **7-508-395** | **90-190** | **>0.5** | **-** | **-** | **-** |
| **7-508-16** | **130-150** | **>0.5** | **-** | **-** | **-** |
| **7-508-39** | **130-150** | **>0.5** | **-** | **-** | **-** |
| **7-508-68** | **130-150** | **>0.5** | **-** | **-** | **-** |
| **7-508-104** | **130-150** | **>0.5** | **-** | **-** | **-** |
| **7-508-415** | **130-150** | **>0.5** | **-** | **-** | **-** |
| **7-508-466** | **370-510** | **>0.5** | **>0.5** | **-** | **-** |
| **12-28-1** | **490-510** | **>0.5** | **>0.5** | **-** | **-** |
| **7-73-121** | **490-510** | **>0.5** | **>0.5** | **-** | **-** |
| **7-508-201** | **490-510** | **>0.5** | **>0.5** | **-** | **-** |
| **7-508-669** | **490-510** | **>0.5** | **>0.5** | **-** | **-** |
| **7-508-362** | **N/A** | **>0.5** | **>0.5** | **>0.5** | **>0.5** |
| **7-508-94** | **N/A** | **>0.5** | **>0.5** | **-** | **-** |
| **7-508-411** | **N/A** | **>0.5** | **>0.5** | **-** | **-** |
| **7-508-478** | **N/A** | **>0.5** | **>0.5** | **-** | **-** |
| **7-508-568** | **N/A** | **>0.5** | **>0.5** | **-** | **-** |
| **7-509-2** | **N/A** | **>0.5** | **>0.5** | **-** | **-** |
| **7-508-73** | **N/A** | **>0.5** | **-** | **-** | **-** |
| **7-508-19** | **N/A** | **>0.5** | **-** | **-** | **-** |
| **7-508-165** | **N/A** | **>0.5** | **-** | **-** | **-** |
| **7-508-254** | **N/A** | **>0.5** | **-** | **-** | **-** |
| **7-508-223** | **N/A** | **>0.5** | **-** | **-** | **-** |
| **7-508-315** | **N/A** | **>0.5** | **-** | **-** | **-** |
| **7-508-528** | **N/A** | **>0.5** | **-** | **-** | **-** |
| **7-508-592** | **N/A** | **>0.5** | **-** | **-** | **-** |
| **7-512-9** | **N/A** | **>0.5** | **-** | **-** | **-** |
| **7-508-143** | **N/A** | **-** | **0.1 - 0.5** | **-** | **-** |
| **7-508-198** | **N/A** | **0.1 - 0.5** | **0.1 - 0.5** | **-** | **-** |
| **7-508-380** | **N/A** | **0.1 - 0.5** | **>0.5** | **-** | **-** |
| **12-28-4** | **N/A** | **-** | **-** | **-** | **-** |
| **7-73-57** | **N/A** | **-** | **-** | **-** | **-** |
| **7-508-448** | **N/A** | **-** | **-** | **-** | **-** |
| **7-508-573** | **N/A** | **-** | **-** | **-** | **-** |
| **7-508-646** | **N/A** | **-** | **-** | **-** | **-** |
| **7-508-680** | **N/A** | **-** | **-** | **-** | **-** |
| **7-508-694** | **N/A** | **-** | **-** | **-** | **-** |

**Extended Data Table 1. Screening of a Panel of SARS-CoV MAbs for Cross-Binding Activity**

A panel of 36 previously generated frozen hybridomas of anti-SARS-CoV MAbs were recovered and scaled up. Hybridoma supernatants were screened for reactivity to the SARS-CoV-2 S protein. Binding for each truncation is reported as absorbance at 450 nm for hybridoma supernatant. Positive cell clones were selected for antibody sequencing.

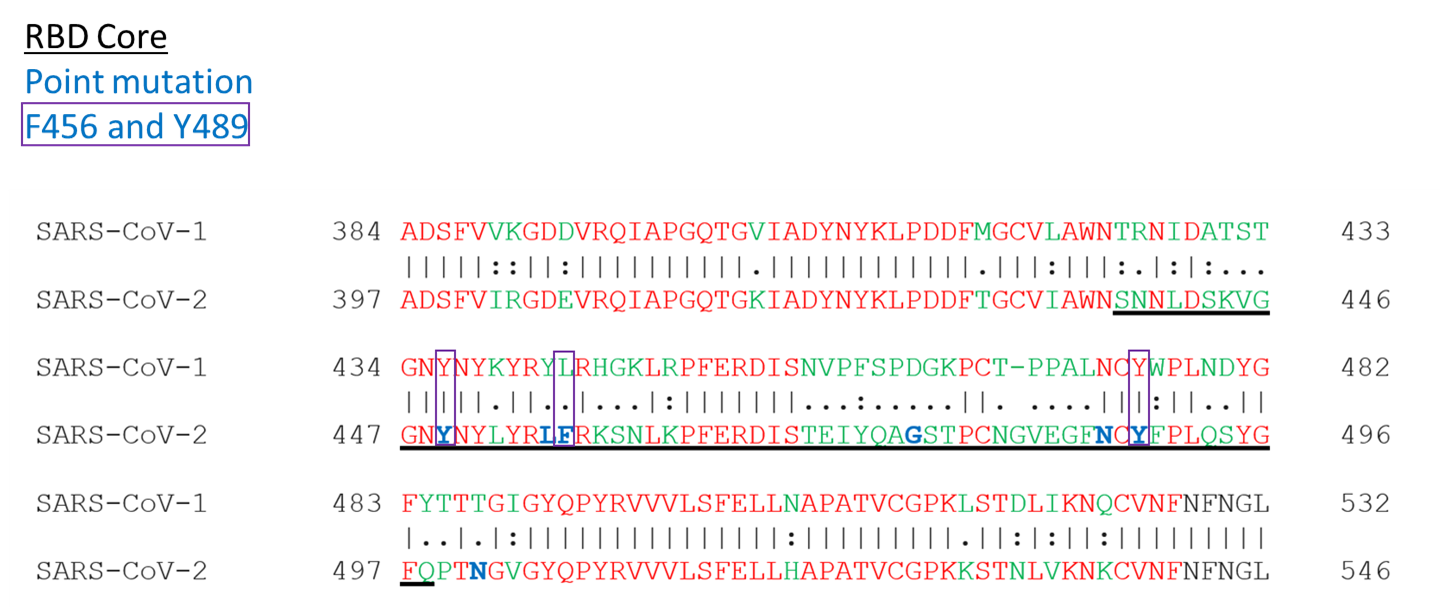
**Extended Data Figure 1. Sequence alignment of SARS-CoV and SARS-CoV-2 S protein RBDs**

Sequence alignment show the identified critical residues (Y449, F456, and Y489) reside within the core domain of the SARS-CoV and SARS-CoV-2 S protein RBD.

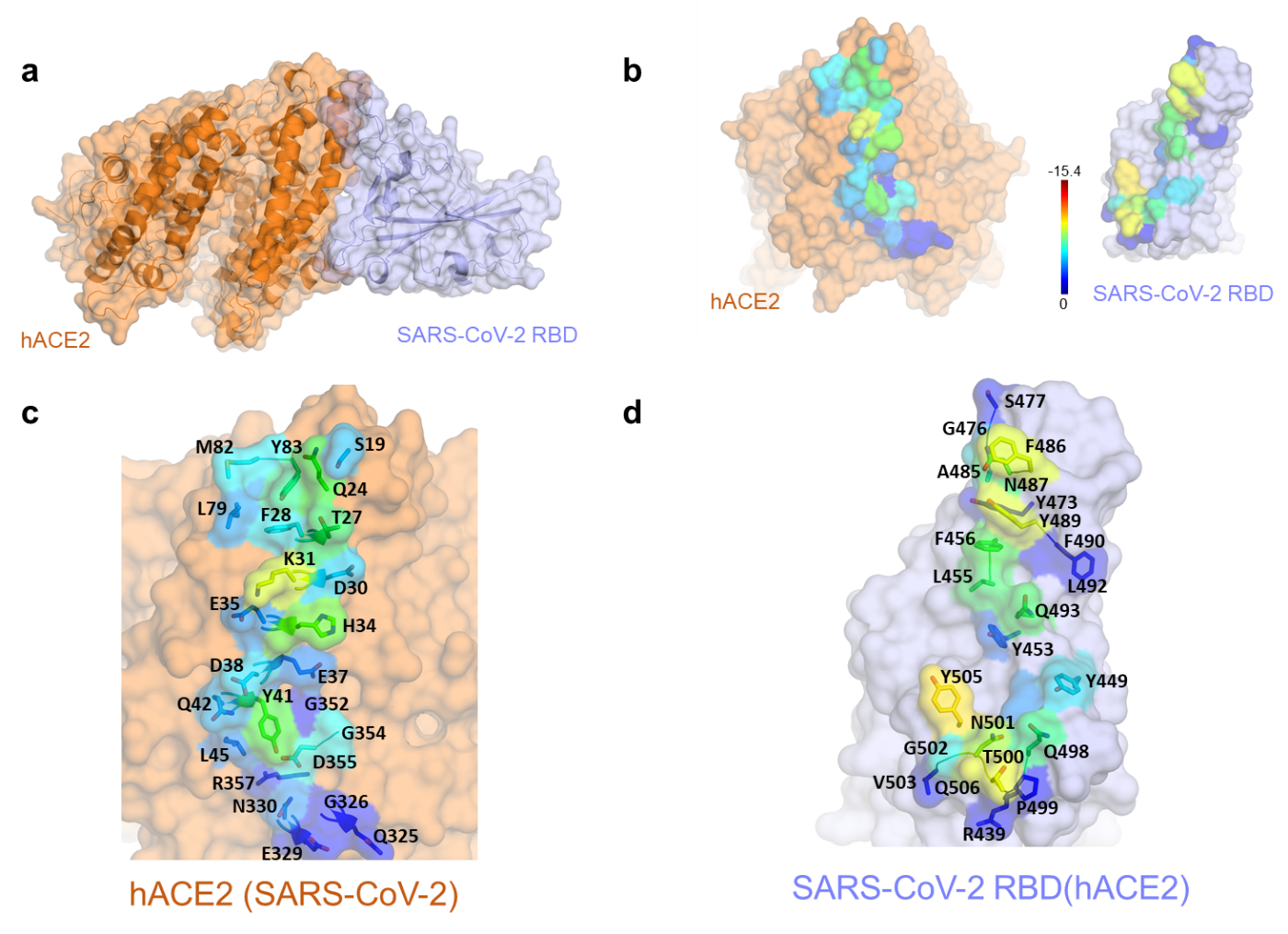
**Extended Data Figure 2. hACE2 - SARS-CoV-2 RBD complex.**

MAb362 is predicted to have an extensive complementary interface with SARS-CoV-2 RBD that extensively overlaps with the hACE2 binding interface. (**a**) Surface representation of the complex, hACE2 in orange, with SARS-CoV-2 RBD in violet. **(b)** The complex is separated and rotated by ~45° to highlight the extensive van der Waals buried surface areas on each protein, the spectrum of color coding represents the extent of the predicted van der Wall contact, with red being the most extensive contact and dark blue the least. **(c)** Detailed van der Waals of hACE2. **(d)** The binding interface on SARS-CoV-2 RBD with hACE2. Residues names shown in red are those that differ in sequence between SARS-CoV and SARS-CoV-2.

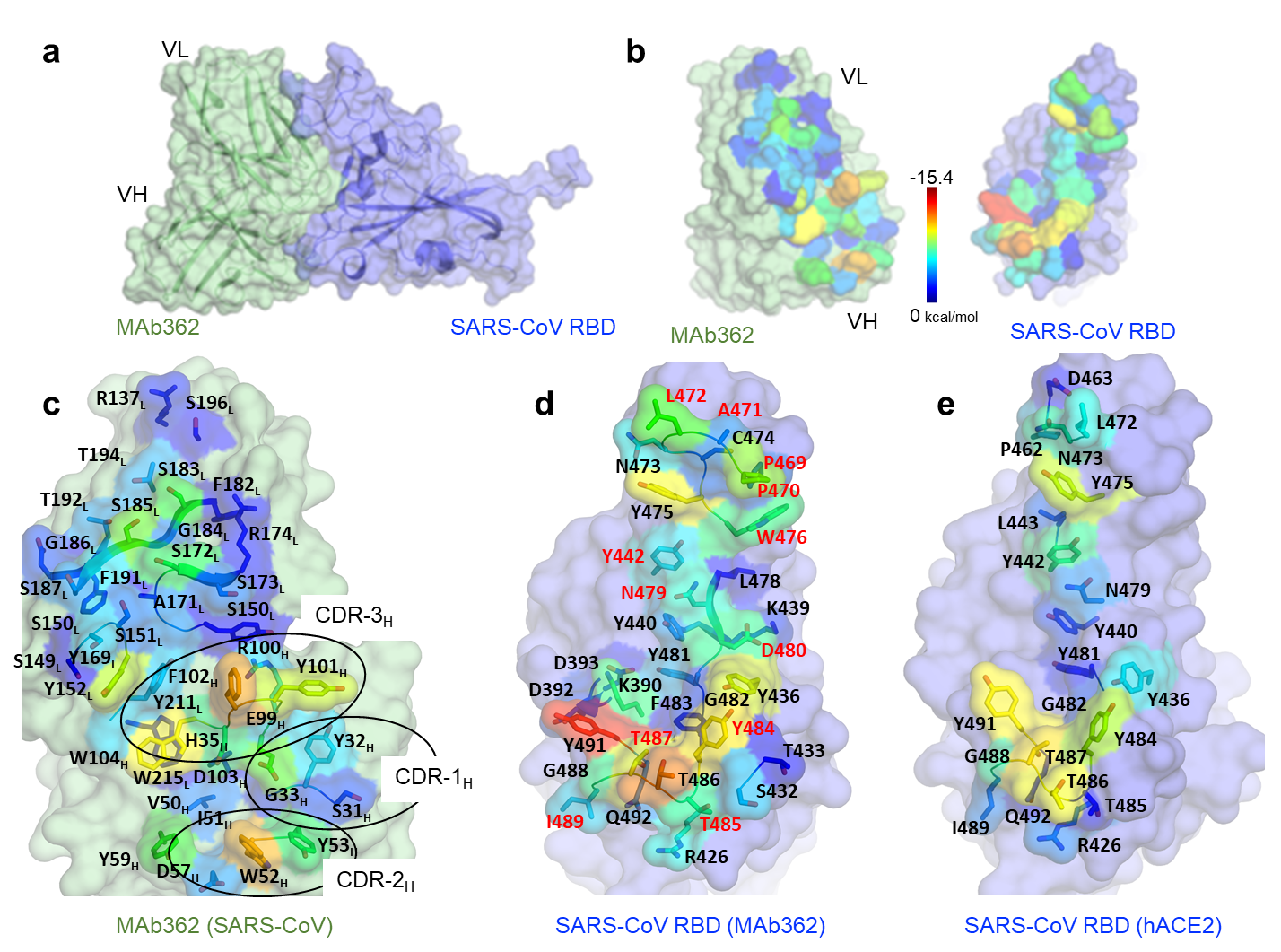
**Extended Figure 3. MAb362 - SARS-CoV RBD complex**

MAb362 is predicted to have an extensive complementary interface with SARS-CoV RBD that extensively overlaps with the hACE2 binding interface. (**a**) Surface representation of the complex, MAb362 in green, with SARS-CoV RBD in violet. **(b)** The complex is separated and rotated by ~45° to highlight the extensive van der Waals buried surface areas on each protein, the spectrum of color coding represents the extent of the predicted van der Wall contact, with red being the most extensive contact and dark blue the least. **(c)** Detailed van der Waals of MAb362, residues from all the CDR’s from both heavy and light chains pack against the SARS-CoV RBD. **(d)** The binding interface on SARS-CoV RBD with MAb362. **(e)** The binding interface on SARS-CoV RBD with hACE2. Residues names shown in red are those that differ in sequence between SARS-CoV and SARS-CoV-2.

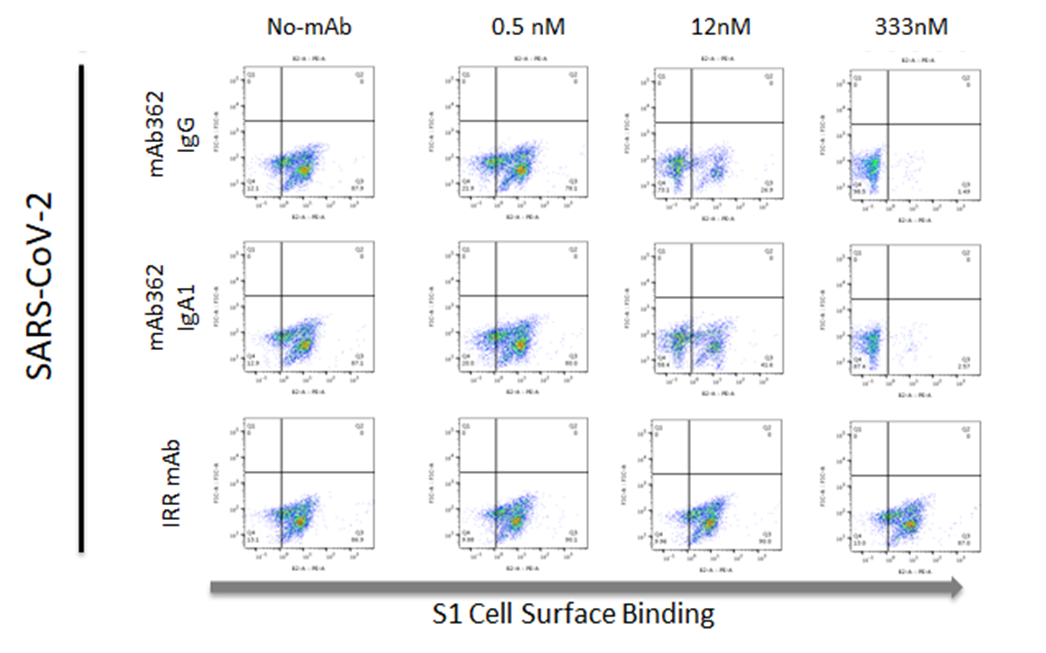
**Extended Data Figure 4. MAb362 blocks SARS-CoV and SARS-CoV-2 binding to hACE2 receptor**

Flow cytometry of MAb362 IgG and IgA1 blocking binding of S_1_ truncations of the SARS-CoV (S_1-590_) and SARS-CoV-2 (S_1-604_) binding to cell surface receptor hACE2 on Vero E6 cells.
